# Targeting the ClpP-αSynuclein Interaction with a Decoy Peptide to Mitigate Neuropathology in Parkinson’s Disease Models

**DOI:** 10.1101/2025.02.16.638540

**Authors:** Di Hu, Xiaoyan Sun, Xin Qi

**Affiliations:** Department of Physiology & Biophysics, Case Western Reserve University School of Medicine, Cleveland, OH 44106, USA; Center for Mitochondrial Research and Therapeutics, Case Western Reserve University School of Medicine, Cleveland, OH 44106, USA

## Abstract

Parkinson’s disease (PD), the most prevalent neurodegenerative movement disorder, is characterized by the progressive loss of dopaminergic (DA) neurons and the accumulation of α-synuclein (αSyn)-rich inclusions. Despite advances in understanding PD pathophysiology, disease-modifying therapies remain elusive, underscoring gaps in our knowledge of its underlying mechanisms. Mitochondria are key targets of αSyn toxicity, and growing evidence suggests that αSyn-mitochondrial interactions contribute to PD progression. Our recent findings identify mitochondrial protease ClpP as a crucial regulator of αSyn pathology, with pathological αSyn binding to and impairing ClpP function, thereby exacerbating mitochondrial impairment and neurodegeneration. To disrupt this deleterious interaction, we developed a decoy peptide, CS2, which directly binds to the non-amyloid-β component (NAC) domain of αSyn, preventing its association with ClpP. CS2 treatment effectively mitigated αSyn toxicity in an αSyn-stable neuronal cell line, primary cortical neurons inoculated with αSyn pre-formed fibrils (PFFs), and DA neurons derived from PD patient-induced pluripotent stem cells (iPSCs). Notably, subcutaneous administration of CS2 in transgenic mThy1-hSNCA PD mice rescued cognitive and motor deficits while reducing αSyn aggregation and neuropathology. These findings establish the ClpP-αSyn interaction as a druggable target in PD and position CS2 as a promising therapeutic candidate for PD and other αSyn-associated neurodegenerative disorders.

## INTRODUCTION

Parkinson’s disease (PD) is the most common neurodegenerative movement disorder, affecting up to 2% of individuals aged 60 and older^1^. It is characterized by the progressive loss of pigmented dopaminergic (DA) neurons in the substantia nigra (SN) of the midbrain and the accumulation of α- synuclein (αSyn)-containing cytoplasmic inclusions known as Lewy bodies^2^. Despite significant advances in understanding PD pathophysiology, no effective strategies have been established to slow disease progression. This persistent therapeutic gap underscores the need for a deeper understanding of the underlying disease mechanisms.

Mitochondrial dysfunction has long been recognized as a key pathological hallmark of PD. Substantial evidence indicates that mitochondria are primary targets of αSyn toxicity, and the interplay between αSyn and mitochondria may play a causal role in PD neuropathology^3, 4^. αSyn contains a cryptic mitochondrial targeting sequence and is enriched in mitochondria within vulnerable brain regions in PD^5^. Mitochondrial accumulation of αSyn induces bioenergetic deficits and increases reactive oxygen species (ROS) production, exacerbating αSyn toxicity, DA neuron degeneration, and associated cognitive and motor deficits^6, 7^. Within mitochondria, disturbances in protein homeostasis due to compromised quality control mechanisms lead to mitochondrial protein overload and stress, ultimately resulting in bioenergetic failure, protein aggregation, and DA neuron degeneration in various PD models^8, 9^. This underscores the pathogenic significance of mitochondrial proteotoxicity.

The mitochondrial matrix protease ClpP plays a crucial role in maintaining mitochondrial proteostasis by processing unfolded or misfolded proteins for degradation, functioning similarly to a proteasome^10, 11^. Our recent findings revealed, for the first time, an interdependence between pathological αSyn and ClpP. αSyn directly binds to ClpP and inhibits its peptidase activity, leading to the accumulation of unfolded mitochondrial proteins and subsequent bioenergetic deficits^12^. Conversely, compensating for ClpP loss has been shown to counteract αSyn-induced mitochondrial and neuronal degeneration in DA neurons derived from PD patient-induced pluripotent stem cells (iPSCs) and to ameliorate αSyn aggregation and motor deficits in αSyn A53T PD mice^12^. These findings suggest that the interaction between αSyn and ClpP may be a critical driver of neurodegeneration in PD and other synucleinopathies.

In this study, to investigate the causal role of the αSyn-ClpP interaction in PD, we developed a decoy peptide, CS2, designed to inhibit this binding. Validation studies demonstrated that CS2 directly binds to the non-amyloid-β component (NAC) domain of αSyn, thereby disrupting the interaction between ClpP and αSyn both *in vitro* and *in vivo*. As a result, CS2 treatment significantly restored ClpP protein expression in αSyn stable cell cultures, primary cortical neurons inoculated with αSyn pre-formed fibrils (PFFs), and DA neurons derived from PD patient iPSCs. In αSyn transgenic mice (mThy1-hSNCA), subcutaneous treatment with CS2 ameliorated cognitive and motor deficits while reducing neuroinflammation and αSyn aggregates in the substantia nigra. These findings demonstrate the ClpP-αSyn interaction as a druggable target in αSyn-associated PD and suggest that CS2-like small peptides could serve as promising therapeutic leads for PD and other neurological disorders characterized by αSyn aggregation.

## RESULTS

### ClpP regulates αSyn tetramer, aggregation and propagation

Our previous findings demonstrated that upregulation of ClpP abolished the expression of phosphorylated serine 129 α-synuclein (pS129-αSyn), a pathological species seen in PD patient, in the substantia nigra (SN) of transgenic A53T-αSyn mice. These results suggest that ClpP play an important role in modulating αSyn pathology^12^. Native αSyn primarily exists as a tetramer that resists aggregation, whereas pathogenic stimuli, such as mitochondrial or lysosomal dysfunction, can destabilize this tetrameric structure, promoting its transition to monomeric αSyn, which is prone to aggregation^13–15^. Therefore, we first investigated whether ClpP attenuates αSyn pathology by stabilizing its tetrameric state. In dopaminergic SH-SY5Y neuronal cells, ClpP overexpression or knockdown significantly increased or decreased the αSyn tetramer-to-monomer ratio, respectively, despite having minimal effect on total αSyn protein levels (Fig. 1A-C). Consistent with previous studies^16^, the αSyn tetramer-to-monomer ratio was significantly reduced in HEK293T cells overexpressing A53T-αSyn compared to those expressing wild-type (WT)-αSyn (Fig. S1A). Notably, overexpression of ClpP in HEK293T cells expressing WT-αSyn significantly elevated the αSyn tetramer-to-monomer ratio (Fig. S1B). Intriguingly, co-overexpression of the proteolytically inactive ClpP mutant (ClpP-S153A) significantly decreased the WT-αSyn tetramer-to-monomer ratio (Fig. S1C). Collectively, these results suggest that ClpP modulates αSyn tetramerization, potentially through its proteolytic activity.

**Figure 1.**
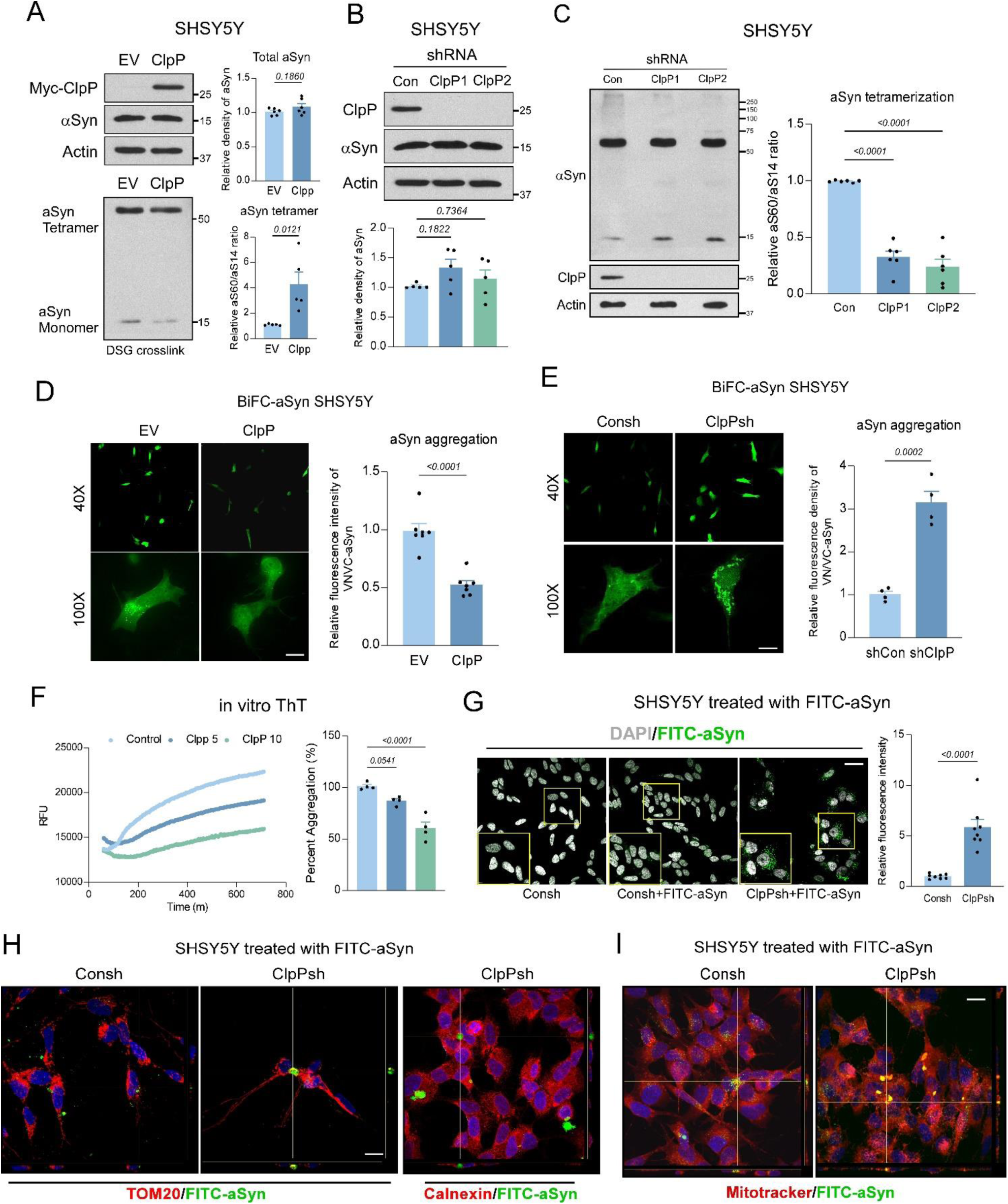
ClpP modulates α-Syn tetramer and aggregates. (A) Top: total lysates were harvested from SH-SY5Y cells overexpressing control empty vector (EV) or Myc-tagged ClpP (Myc-ClpP) and subjected for western blot (WB) analyses; bottom: SH-SY5Y cells overexpressing EV or Myc-ClpP were subjected for intact cell crosslinking by DSG (see methods), and WB to examine the level of different α-Syn species. Shown are the representative blots of 6 independent repeats. Histograms: the relative expression of total endogenous α-Syn or the ratio of α-Syn tetramer to monomer. (B) Total lysates were harvested from SH-SY5Y cells stably expressing control shRNA (Con) or different shRNA against ClpP (ClpP1/2) and subjected for WB analyses. Shown are the representative blots of 5 independent repeats Histogram: the relative expression of total endogenous α-Syn. (C) SH-SY5Y cells expressing Con or ClpP1/2 shRNA were subjected for intact cell crosslinking by DSG, and WB analyses. Shown are the representative blots of 6 independent experiments. Histogram: the relative ratio of α-Syn tetramer to monomer. (D) SH-SY5Y cells overexpressing EV or ClpP were transfected with vectors expressing Venus-N-α-Syn (VN) and α-Syn-C-Venus (VC) to monitor α-Syn oligomerization. Shown are the representative images at different magnifications of 8 independent experiments. Scale bar=10µm. Histogram: relative fluorescence intensity of α-Syn BiFC. (E) SH-SY5Y expressing Con or ClpP shRNA were transfected with VN and VC vectors to monitor α-Syn oligomerization. Shown are the representative images at different magnifications of 4 independent experiments. Scale bar=10µm. Histogram: relative fluorescence intensity of α-Syn BiFC. (F) α-Syn (5µM) aggregation in the presence and absence of ClpP (5/10µM) was monitored by thioflavin T (ThT) fluorescence. Shown are the representative curves of 4 independent experiments. Histogram: relative α-Syn aggregates forms in the indicated conditions. (G) SH-SY5Y cells expressing Con (Consh) or ClpP (ClpPsh) shRNA were treated with FITC-labeled α-Syn monomer for 24h. Shown are the representative images of 8 independent experiments. Scale bar=30µm. Enlarged area were circled in yellow. Histogram: relative fluorescence intensity of FITC-α-Syn. Consh and ClpPsh expressing SH-SY5Y cells were treated with FITC-α-Syn monomer for 24h and subjected for (H) immunostaining using anti-Tom20 or anti-Calnexin antibody or (I) Mitotracker dye staining. Scale bar=20 µm. Shown are the representative z-stack images of 3 independent experiments. All data are expressed as the mean ± SEM. Data are compared with one-way ANOVA with Tukey’s *post-hoc* test in B, C, F. Unpaired student *t*-test is used in A, D, E, G.

We next examined whether ClpP modulates αSyn aggregation and propagation, processes that are stimulated upon disruption of αSyn tetramers. αSyn oligomers can be quantitatively visualized using constructs expressing bimolecular fluorescence complementation (BiFC) VN/VC-αSyn (Venus-N-αSyn and αSyn-C-Venus)^17^. In SH-SY5Y cells, ClpP overexpression significantly reduced VN/VC-αSyn fluorescence intensity (Fig. 1D), indicating a decrease in αSyn oligomerization. Conversely, ClpP knockdown led to a substantial increase in αSyn oligomers in SH-SY5Y cells (Fig. 1E). These findings suggest that ClpP regulates αSyn oligomerization. To further investigate whether ClpP modulates αSyn aggregation, we employed an *in vitro* αSyn PFF-seeded aggregation assay, which can be quantified using Thioflavin T (ThT) fluorescence^18, 19^. Incubation with recombinant ClpP significantly reduced PFF-ThT fluorescence intensity in a dose-dependent manner, indicating that ClpP directly modulates αSyn aggregation (Fig. 1F). To assess ClpP’s role in αSyn propagation, we treated SH-SY5Y cells expressing either control or ClpP shRNA with FITC-conjugated αSyn. ClpP deficiency resulted in substantial accumulation and aggregation of FITC-conjugated αSyn in SH-SY5Y cells (Fig. 1G), suggesting that ClpP loss promotes αSyn spreading and accumulation. Recent Cryo-ET and TEM studies implicate mitochondria as a preferential platform for αSyn accumulation and aggregation^20^. Consistent with this evidence, FITC-αSyn formed detergent-resistant puncta that colocalized with the mitochondrial marker TOM20 and Mitotracker Red but not with the ER protein calnexin in ClpP-deficient cells (Fig. 1H, I). Together, these results indicate that ClpP negatively regulates αSyn oligomerization, aggregation, and propagation.

### ClpP interacts with αSyn NAC domain

To gain mechanistic insights into the interaction between ClpP and αSyn, we examined which protein domain of αSyn binds to ClpP and contributes to proteolytic impairment. Previous structural analyses using X-ray crystallography and Cryo-EM revealed that ClpP consists of a series of highly conserved α-helices connected by flexible motifs^21^. In contrast, αSyn is primarily composed of three peptide domains: the N-terminal domain, the NAC domain, and the C-terminal domain. Among these, the NAC domain is essential for amyloid core formation during αSyn aggregation^22^. To determine the domain responsible for ClpP binding, we performed co-immunoprecipitation (Co-IP) in HEK293T cells overexpressing full-length ClpP along with truncated αSyn mutants lacking either the N-terminal (ΔN), C-terminal (ΔC), or NAC domain (ΔNAC). While WT, ΔN, and ΔC αSyn bound to ClpP, the ΔNAC mutant abolished the interaction (Fig. 2A). Consistently, an in vitro Co-IP assay further confirmed the absence of direct binding between ΔNAC αSyn and ClpP recombinant protein (Fig. 2B), suggesting that ClpP specifically interacts with the αSyn NAC domain. To assess whether the NAC domain is required for αSyn-induced ClpP downregulation, we examined ClpP protein levels in HEK293T cells overexpressing WT or truncated αSyn mutants. Overexpression of WT, ΔN, or ΔC αSyn significantly decreased ClpP expression, whereas ΔNAC αSyn had minimal effect (Fig. 2C). Functionally, WT and ΔC αSyn substantially impaired ClpP peptidase activity, whereas ΔNAC αSyn exhibited minimal effects (Fig. 2D). In summary, these findings suggest that the αSyn NAC domain is required for its interaction with and modulation of ClpP.

**Figure 2.**
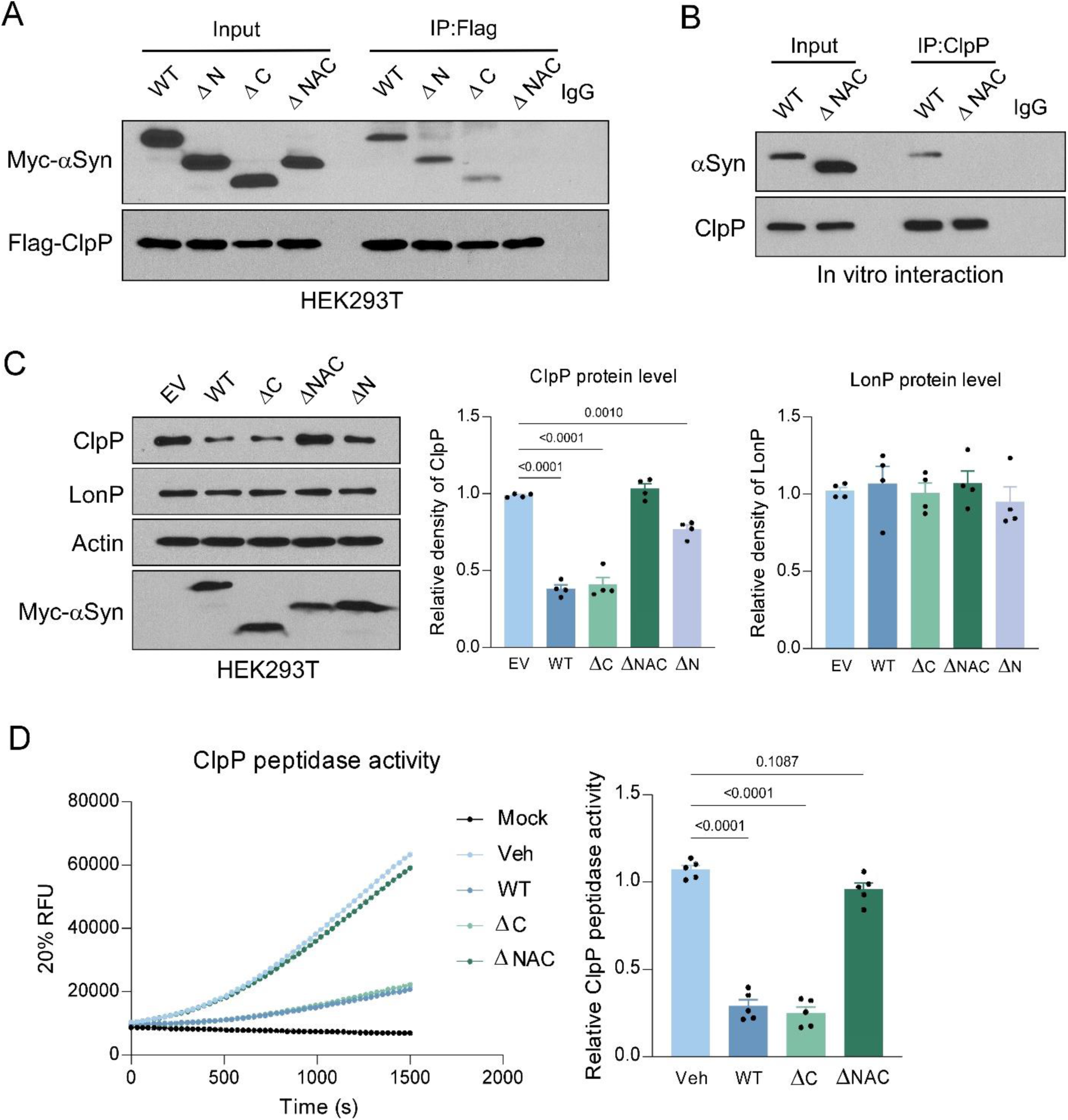
α-Syn NAC domain impairs ClpP. (A) Total lysates were harvested from HEK293T cells overexpressing Myc-tagged wild-type (WT), N-terminal truncated (ΔN), C-terminal truncated (ΔC), or NAC-domain truncated (ΔNAC) α-Syn with Flag-tagged ClpP and subjected for immunoprecipitation using anti-Flag antibody, followed by western blot (WB) analyses. Shown are the representative blots of 3 independent experiments. (B) Recombinant ClpP was incubated with either WT or ΔNAC α-Syn in vitro for 12h, and pulled down using anti-ClpP antibody, followed by WB analyses. Show are the representative blots of 3 independent experiments. (C) Total lysates were harvested from HEK293T cells overexpressing Myc-tagged WT, ΔN, ΔC, or ΔNAC α-Syn and subjected for WB analyses. Shown are the representative blots of 4 independent experiments. Histogram: relative expression of ClpP and LonP. (D) In vitro ClpP peptidase activity was measured in the absence or presence WT, ΔC, or ΔNAC α-Syn. The fluorescence intensity of ac-WLA-AMC (50 µM), a fluorogenic substrate of ClpP, was measured up to 30 min immediately after the addition. Shown are the representative RFU measurement of 5 independent experiments. Histogram: relative ClpP peptidase activity. All data are expressed as the mean ± SEM. Data are compared with one-way ANOVA with Tukey’s *post-hoc* test.

### Decoy peptide CS2 disrupts ClpP/αSyn interaction

We next assessed the translational potential of disrupting the interaction between αSyn and ClpP in PD. Previously, we designed several peptide inhibitors that effectively rescued mitochondrial dysfunction and prevented neuropathological progression in models of neurodegenerative diseases, including Alzheimer’s disease (AD) and Huntington’s disease (HD), by targeting specific protein-protein interactions^23–26^. Leveraging this approach, we designed and developed two peptides, CS1 and CS2, derived from homologous sequences within αSyn and ClpP, respectively (Fig. 3A). Both peptides were conjugated to an HIV TAT carrier at the N-terminus to enhance transmembrane delivery and facilitate blood-brain barrier penetration ^24^. To determine whether CS1 or CS2 could block the interaction between αSyn and ClpP in vitro, we pre-incubated ClpP with either peptide and assessed binding. While CS1 had no effect, CS2 significantly diminished αSyn-ClpP interaction (Fig. 3B), indicating that CS2, rather than CS1, effectively interferes with the binding. We then tested CS2’s efficacy in cell culture. In SH-SY5Y cells, intracellular fluorescence intensity of FITC-conjugated CS2 peptide peaked at approximately 30 minutes, demonstrating efficient transmembrane delivery (Fig. S2). Given that CS2 is derived from a homologous ClpP sequence, we hypothesized that it directly binds to αSyn. To test this, we incubated biotin-conjugated CS2 peptide with total protein lysates containing overexpressed ClpP or αSyn, followed by streptavidin pull-down. CS2 selectively interacted with αSyn but not with ClpP or LonP (a mitochondrial matrix protease) (Fig. 3C). Furthermore, CS2 treatment abolished ClpP’s interaction with WT, ΔC, and ΔN αSyn in HEK293T cells (Fig. 3D-F), confirming its ability to disrupt αSyn-ClpP binding. We further examined the effects of CS2 treatment on ClpP expression. In both SH-SY5Y and HEK293T cells, CS2 treatment significantly restored ClpP protein levels, which had been reduced by WT- or A53T-αSyn expression (Fig. 3G, H). Similarly, in Tet-on inducible αSyn-expressing SH-SY5Y cells, CS2 treatment led to a substantial increase in ClpP protein expression compared to the TAT-treated control group (Fig. 3I). Collectively, these findings indicate that CS2 efficiently disrupts the αSyn-ClpP interaction, thereby restoring ClpP expression.

**Figure 3.**
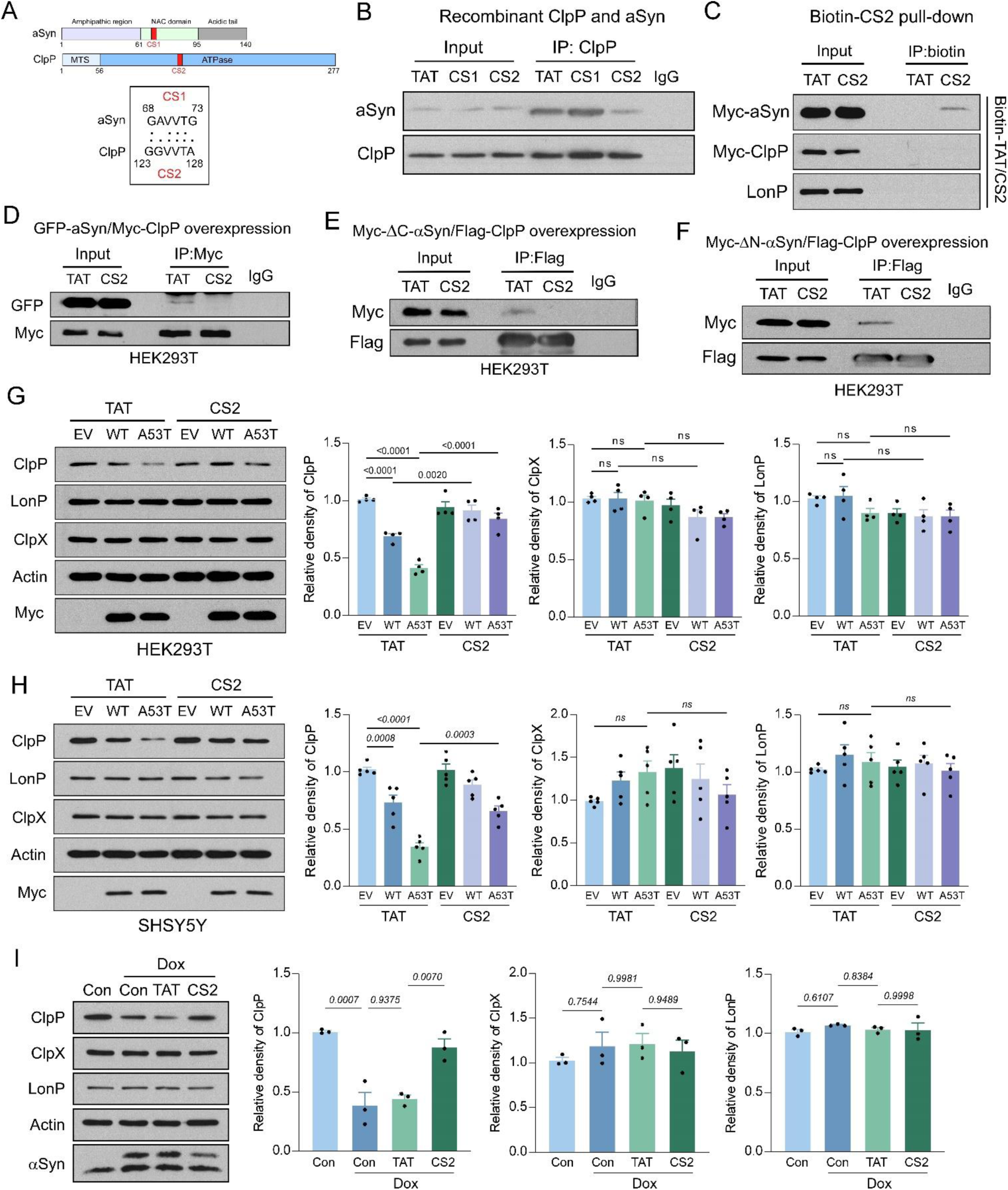
Rationally design peptide disrupts the interaction between ClpP and α-Syn. (A) Domain maps of human ClpP and α-Syn. Peptides CS1 and CS2 are derived from the highly conserved peptide sequence between ClpP and α-Syn as indicated in red. (B) Recombinant ClpP was pre-incubated with control peptide TAT, or CS1/CS2 for 30 min, before adding α-Syn, and subjected for immunoprecipitation using anti-ClpP antibody, followed by western blot (WB) analyses. Shown are representative blots of 3 independent experiments. (C) Total lysates were harvested from HEK293T cells overexpressing Myc-tagged α-Syn or ClpP and incubated with biotin-labeled TAT or CS2 peptides for 12h in vitro. The biotin-TAT or CS2 was pulled down by using streptavidin-beads and subjected for WB analyses. Shown are the representative blots of 3 independent experiments. (D) Total lysates were harvested from HEK239T cells expressing GFP-tagged α-Syn and Myc-tagged ClpP upon TAT or CS2 treatment, and subjected for immunoprecipitation using anti-Myc antibody, followed by WB analyses. Shown are representative blots of 3 independent experiments. Total lysates were harvested from HEK293T cells overexpressing Flag-tagged ClpP with (E) Myc-tagged ΔC-α-Syn or (F) Myc-tagged ΔN- α-Syn, and subjected for immunoprecipitation using anti-flag antibody, followed by WB analyses. Shown are representative blots of 3 independent experiment. (G) HEK293T cells and (H) SH-SY5Y cells were pre-treated with TAT or CS2 30 min before transfection of EV or vectors expressing WT or A53T α-Syn. 48h after transfection, total lysates were harvested and subjected for WB analyses. Shown are the representative blots of (G) 4 and (H) 5 independent experiments. Histogram: relative expression of the indicated proteins. (I) Tet-on inducible SHSY5Y cells were pre-treatment with TAT or CS2 30 min before adding Dox to induce α-Syn overexpression. 72h after adding Dox, total lysates were harvested and subjected for WB analyses. Shown are the representative blots of 3 independent experiments. Histogram: relative expression of the indicated proteins. All data are expressed as the mean ± SEM. Data are compared with one-way ANOVA with Tukey’s *post-hoc* test.

### CS2 binds to αSyn and attenuate its cytotoxicity

To better understand whether and how CS2 attenuates αSyn-induced cytotoxicity, we investigated its binding affinity to αSyn using microscale thermophoresis (MST). Given that CS2 directly binds to αSyn (Fig. 3C), we quantified this interaction and found that CS2 exhibited a dissociation constant (Kd) of 7 µM, whereas the TAT control peptide showed no binding trend toward αSyn (Fig. 4A). Notably, CS2 incubation significantly reduced αSyn-PFF-seeded fibrillization in vitro in a dose-dependent manner (Fig. 4B), indicating that CS2 directly modulates αSyn aggregation. Furthermore, CS2 treatment significantly restored ClpP peptidase activity, which had been impaired by WT- or A53T-αSyn (Fig. 4C, D). Importantly, treatment with TAT or CS2, even at concentrations up to 100 µM, had no observed effect on ClpP peptidase activity, suggesting minimal toxicity (Fig. S3A). Collectively, these in vitro findings highlight the protective role of CS2 against αSyn toxicity.

**Figure 4.**
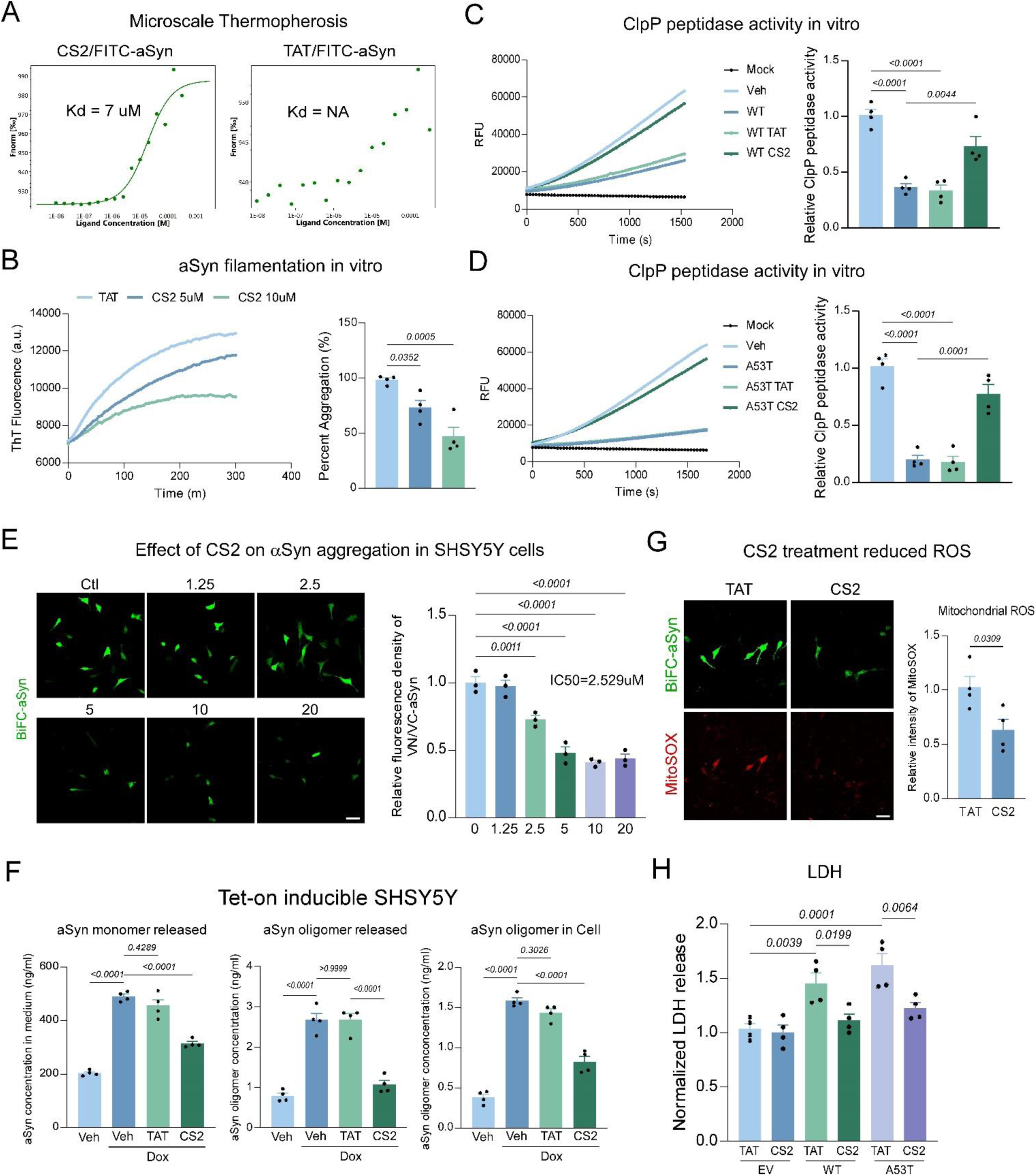
CS2 treatment rescued α-Syn induced toxicity. (A) The binding affinity (Kd = 7µM) of CS2 with α-Syn was determined by MicroScale Thermophoresis (see methods). Shown are representative curve of 3 independent experiments. (B) Effect of TAT or CS2 peptides on α-Syn PFF-seeded fibrillization was examined by dynamic Thioflavin T assay (see methods). Thioflavin T fluorescence intensity was measured every 5 min. Shown are the representative blots of 4 independent experiments. Histogram: relative ThT intensity was normalized to TAT group. ClpP peptidase activity was measured in the presence of (C) WT- or (D) A53T-α-Syn and TAT or CS2 peptides. Shown are the representative RFU measurement of 4 independent experiments. (E) SH-SY5Y cells were pre-treated with CS2 peptides at indicated doses and transfected with vectors expressing VN/VC- α-Syn. The VNVC bimolecular fluorescence was examined 24h after transfection. Shown are the representative images of 3 independent experiments. Scale bar=30µm. Histogram: relative fluorescence intensity of VNVC-α-Syn. (F) SH-SY5Y cells were pre-treated with TAT or CS2 and transfected with VNVC-α-Syn vectors. 24h after transfection, mitochondrial ROS level was measured by MitoSOX staining. Shown are the representative images of 3 independent experiments. Scale bar=30µm. Histogram: relative intensity of mitoSOX. (G) SH-SY5Y cells stably overexpressing EV, WT or A53T α-Syn were treated with TAT or CS2 peptides for 2 days before 12h-serum starvation. Cell death was examined by measuring LDH released in the culture medium. Histogram: normalized level of LDH released. n=4 independent experiments. (H) Tet- on inducible SH-SY5Y cells were pre-treated with TAT or CS2 peptides 30min before adding Dox to induce α-Syn overexpression. 72h after adding Dox, the level of α-Syn monomer in medium, the level of α-Syn oligomer in medium and cytosol were examined by ELISA (see methods). Histogram: relative concentration of α-Syn monomer and oligomers. n=4 independent experiments. All data are expressed as the mean ± SEM. Data are compared with one-way ANOVA with Tukey’s *post-hoc* test. Unpaired student *t*-test is used in F.

Next, we examined the protective effects of CS2 in αSyn-associated PD cell culture models. In SH-SY5H cells expressing VN/VC-αSyn, CS2 treatment led to significant reduction in the fluorescence intensity of BiFC-αSyn with IC50 = 2.5µM, which supports the regulatory effect of CS2 on α-synuclein aggregation. We then tested whether CS2 can mediate α-synuclein propagation. In the Tet-on inducible SH-SY5Y cells, comparing with TAT control peptide, CS2 treatment led to significant decrease in the concentration of extracellular αSyn monomer and oligomers, and intracellular αSyn oligomer, indicating that CS2 can reduce αSyn spreading (Fig. 4F). Given that αSyn-induced ClpP loss caused mitochondrial oxidative stress^12^, we then evaluated the effect of CS2 on the production of mitochondrial ROS. Comparing to TAT peptide, CS2 treatment resulted in significant reduction of the fluorescence intensity of MitoSOX in SH-SY5Y cells expressing BiFC αSyn (Fig. 4G). Furthermore, CS2 treatment mitigated cell death in SH-SY5Y cells expressing either WT- or A53T-α-Syn (Fig. 4H). It is noteworthy that CS2 peptides did not affect cell viability nor ATP production in SH-SY5Y cells (Fig. S3B and C), indicating minor toxicity. In summary, these results indicate that CS2 can attenuate αSyn-induced cytotoxicity.

### CS2 treatment alleviates αSyn-PFF induced neural toxicity

We next evaluated the protective effect of CS2 in αSyn-PFF-inoculated primary cortical neurons, a widely used *ex vivo* model for studying αSyn-induced neurotoxicity. Compared to control neurons, αSyn-PFF inoculation resulted in ClpP co-localization with αSyn, which was significantly diminished upon CS2 treatment (Fig. 5A, B). Additionally, CS2 treatment markedly reduced the fluorescence intensity of pS129-αSyn in PFF-inoculated neurons (Fig. 5C, D), supporting our observation that CS2 mitigates αSyn aggregation and propagation. The interaction of ClpP with αSyn facilitates the transfer of ClpP from detergent-soluble to detergent-insoluble fractions, suggesting ClpP co-aggregation with αSyn both in vitro and in vivo^12^. Consistent with our previous findings, after extracting soluble proteins using Triton X-100 during neuronal fixation, both ClpP and αSyn were detected in αSyn-PFF-inoculated neurons but not in control neurons (Fig. 5E, F). Notably, CS2 treatment significantly reduced ClpP fluorescence intensity in the insoluble fraction (Fig. 5E, F), indicating that CS2 prevents ClpP/αSyn interaction and co-aggregation. To further assess the neuroprotective effects of CS2, we examined synaptic integrity. In line with previous studies^27, 28^, αSyn-PFF inoculation caused synaptic impairment, as evidenced by the uncoupling of pre- and post-synaptic proteins in cortical neurons (Fig. 5G, H). However, CS2 treatment substantially restored synaptic integrity compared to TAT-treated neurons (Fig. 5G, H). Taken together, these findings highlight the protective effects of CS2 against αSyn-PFF-induced neurotoxicity.

**Figure 5.**
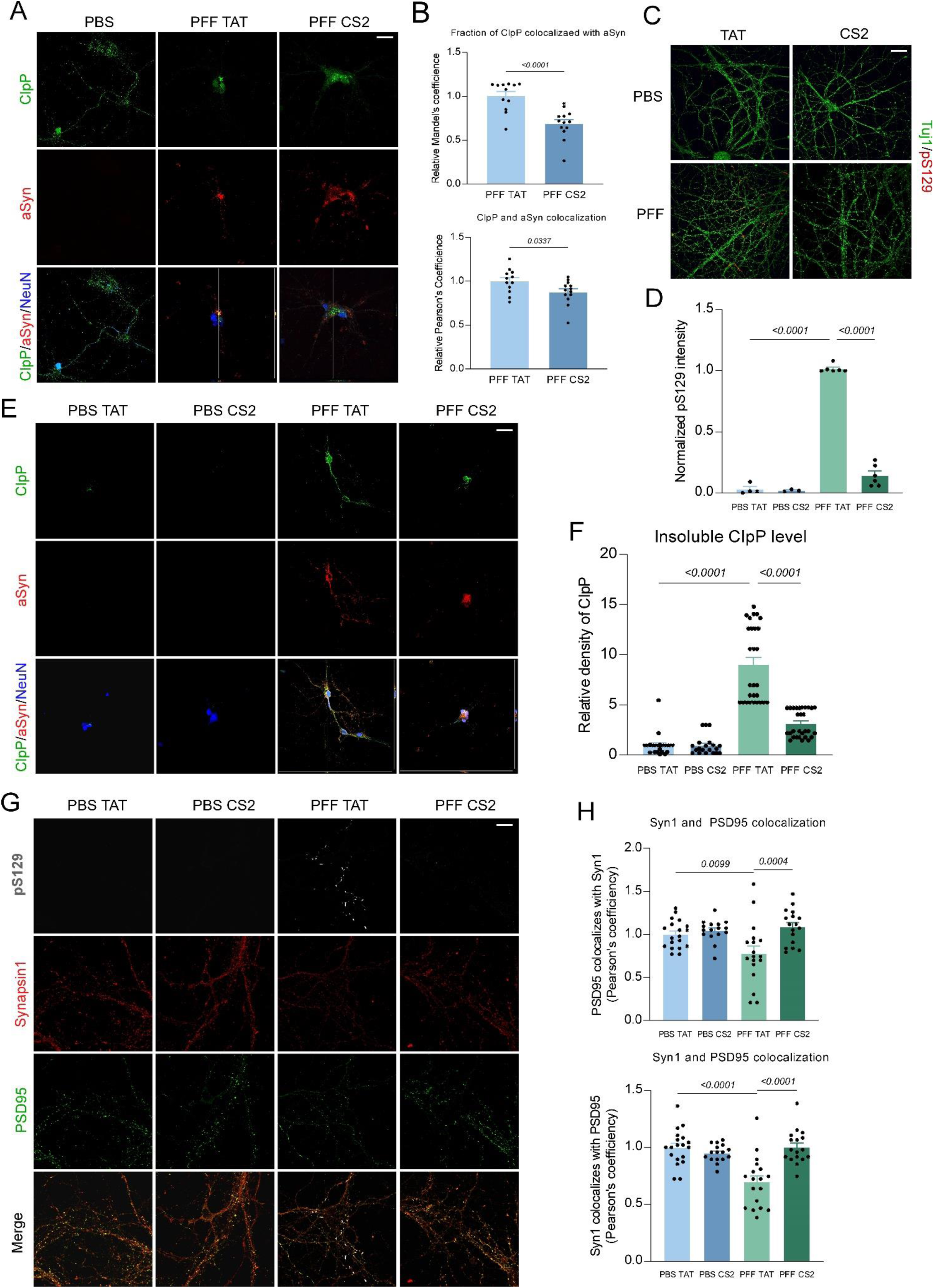
CS2 treatment rescued α-Syn PFF-induced neurotoxicity. Primary cortical neurons were isolated from E18 mouse cortex (see methods) and plated on coverslips (Day 0). Neurons were treated with human α-Syn PFF on Day 5 at 50ng/ml. On Day 7, TAT or CS2 (1µM) was added daily till Day 12. (A) Neurons were fixed in 4% PFA and subjected for immunostaining with anti-ClpP, anti-α-Syn, and anti-NeuN. Shown are the representative Z-stack images of 3 independent experiments. Colocalization of ClpP with α-Syn were examined. N=12 neurons. (B) Histogram: relative Mandel’s co-efficiency and pearson’s co-efficiency for ClpP and α-Syn colocalization. (C) Neurons were fixed in 4% PFA and subjected for immunostaining with anti-Tuj1 and anti-pS129. Shown are the representative images of at least 3 independent experiments. (D) Histogram: related fluorescence intensity of pS129-α-Syn per Tuj1+ area. (E) Neurons were fixed in 4% PFA/4% sucrose/1%Triton-X100 to extract the soluble factions, and subjected for immunostaining with anti-ClpP, anti-α-Syn, and anti-NeuN. Shown are representative Z-stack images of 3 independent experiments. (F) Histogram: relative fluorescence intensity of ClpP in the neuronal insoluble fractions. At least 30 neurons were counted. (G) Neurons were fixed in 4% PFA and subjected for immunostaining with anti-pS129, anti-Synapsin1, and anti-PSD95 antibodies. Shown are the representative images of 3 independent experiments. (H) Histogram: Pearson’s co-efficiency of PSD95/Synapsin1 colocalization. At least 30 areas were counted. Scale bar=30µm. All data are expressed as the mean ± SEM. Data are compared with one-way ANOVA with Tukey’s *post-hoc* test in D, F, H. Unpaired student *t*-test is used in B.

### CS2 treatment is protective in DA neurons derived from PD patients iPSCs

We previously differentiated iPSCs derived from PD patients carrying A53T-αSyn into DA neurons. As a result of toxic A53T-αSyn expression, these patient-derived DA neurons exhibited ClpP downregulation, mitochondrial oxidative stress, and dendritic shortening^12^. Thus, this model provides a valuable platform to validate the protective effects of CS2 against αSyn-induced mitochondrial dysfunction and neuronal damage. Following our established protocol, DA neurons differentiated from A53T or isogenic control iPSCs were treated with TAT or CS2 (1 µM) for five consecutive days^12^. Compared to the isogenic control, pS129-αSyn expression was substantially elevated in tyrosine hydroxylase (TH)-marked DA neurons derived from PD patient iPSCs carrying A53T-αSyn (Fig. 6A, B). Notably, CS2 treatment significantly reduced pS129-αSyn levels in these PD patient DA neurons (Fig. 6A, B). Consistent with previous reports, A53T-αSyn PD patient DA neurons exhibited widespread dendritic loss and synaptic impairments compared to the isogenic control (Fig. 6C-E). CS2 treatment restored the density of both pre- and post-synaptic markers, PSD95 and Synapsin1, in these patient neurons (Fig. 6C-E). These findings further demonstrate the protective role of CS2 against αSyn-induced neurotoxicity.

**Figure 6.**
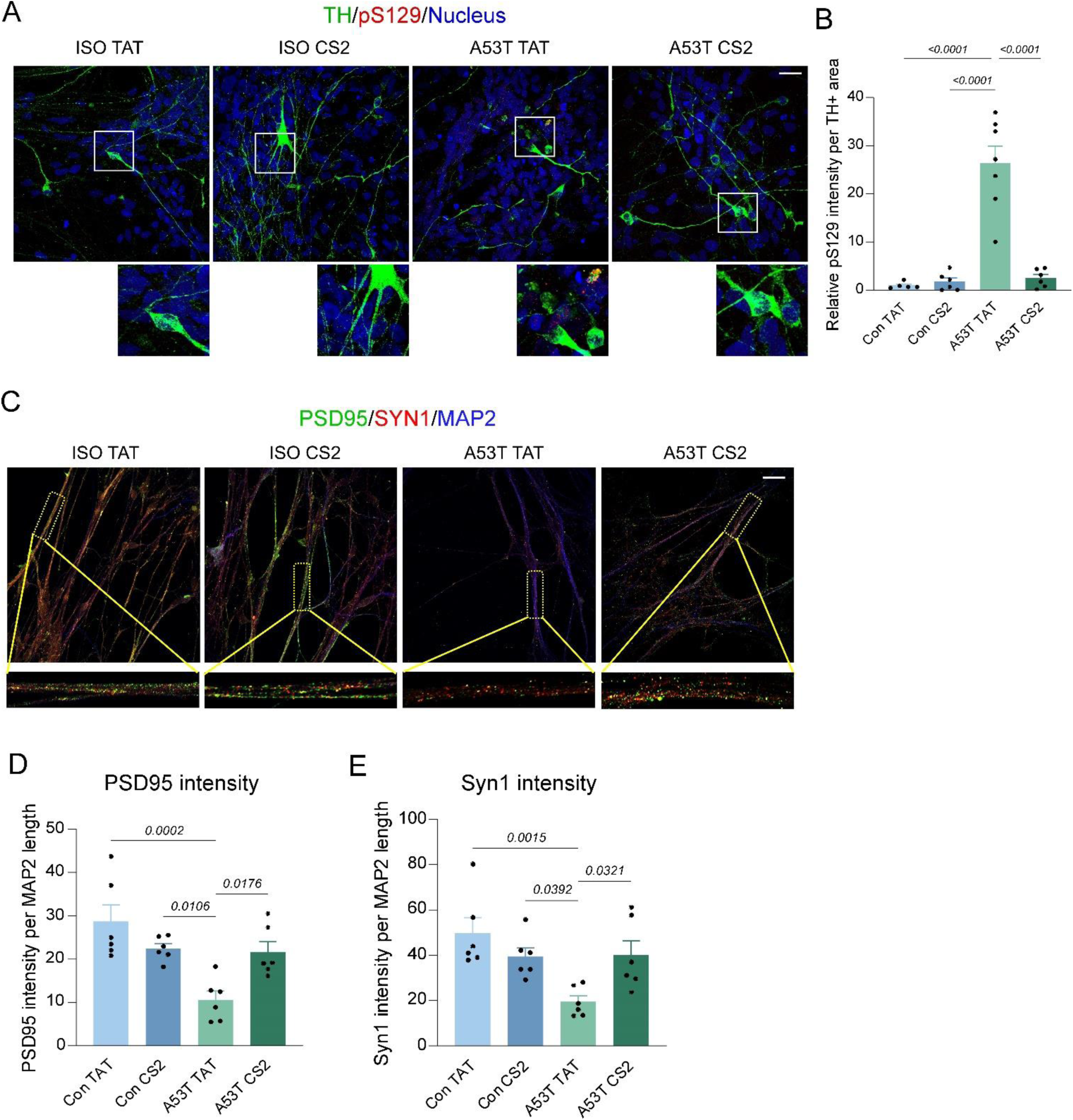
CS2 treatment is protective in DA neurons derived from patient A53T α-Syn iPSCs. A53T α-Syn patient iPSCs (A53T) and its isogenic control iPSCs (ISO) were differentiated into DA neurons (see methods) for 28 days. Neurons were treated with TAT or CS2 for 5 consecutive days starting from Day 28 at 1µM. (A) Neurons were fixed and subjected for immunostaining with anti-TH, anti-pS129 antibodies. Shown are the representative images of at least 5 independent experiments. Enlarged area are circled in white. (B) Histogram: related pS129-α-Syn intensity per TH+ area. (C) Neurons were fixed and subjected for immunostaining with anti-PSD95 and anti-Synapsin1 antibodies. Shown are the representative images of 6 independent experiments. Enlarged neuronal dendrites were circled in yellow. (D) Histogram: relative fluorescence intensity of PSD95 per MAP2 length. (E) Histogram: relative fluorescence intensity of Synapsin1 per MAP2 length. Scale bar=30µm. All data are expressed as the mean ± SEM. Data are compared with one-way ANOVA with Tukey’s *post-hoc* test.

### CS2 treatment alleviates αSyn-associated neuropathology and motor deficits *in vivo*

Next, we examined whether sustained CS2 treatment attenuates αSyn-associated neuropathology *in vivo*. The transgenic mThy1-hSNCA mouse (Line 15), which expresses wild-type human αSyn under the control of the mouse Thy-1 promoter, is widely used to study PD and αSyn aggregation-induced neurodegeneration. In addition to PD-associated neuropathology and motor deficits, mThy1-hSNCA mice exhibit early-stage (4.5 months) cognitive impairments reminiscent of Dementia with Lewy Bodies (DLB)^29^. Accordingly, beginning at 3 months of age, TAT or CS2 peptides (3 mg/kg/day) were administered subcutaneously via an osmotic pump to mThy1-hSNCA mice and their wild-type (WT) littermates. Preliminary evidence from a small-cohort study supports this dosage. We first evaluated cognitive function in the TAT- and CS2-treated mice. At 5 months of age, compared to their WT littermates, mThy1-hSNCA mice exhibited significant deficits in short-term memory and spatial learning, as indicated by a reduced alternation ratio in the Y-maze test (Fig. 7A). Notably, CS2 treatment significantly increased the alternation ratio in mThy1-hSNCA mice (Fig. 7A), suggesting cognitive improvement. We then assessed locomotor activity in mThy1-hSNCA mice at 8 months using an open-field chamber. Compared to WT littermates, mThy1-hSNCA mice were hyperactive, as reflected by increased total traveling distance and vertical activity counts; CS2 treatment normalized these abnormalities (Fig. 7B). Additionally, mThy1-hSNCA mice exhibited impaired motor coordination on the rotarod test, which was ameliorated by sustained CS2 treatment (Fig. 7C). Importantly, CS2 treatment had no observed effect on WT mice, indicating minimal toxicity even after prolonged administration (Fig. 7). Collectively, these results demonstrate that CS2 treatment protects against αSyn-associated cognitive impairments and motor deficits in vivo. We then determined the effect of CS2 on neuropathology in the mThy1-hSNCA mice. Consistent with what we have detected in the hA53T-αSyn expressing mice, the protein level of ClpP was selectively and significantly decreased in the mid-brain of the mThy1-hSNCA mice, comparing with their WT littermates (Fig. 7D and E). CS2 treatment rescued the expression of ClpP, but not other mitochondrial matrix proteins, in the mid-brain of the mThy1-hSNCA mice (Fig. 7D and E). Moreover, the protein level of pS129-αSyn was significantly decreased after sustained treatment of CS2 in the mThy1-hSNCA mice (Fig. 7D and E). CS2 treatment led to significant reduction of αSyn level in both the detergent-soluble and -insoluble fractions (Fig. 7D and E), and mitigated αSyn aggregation in the mid-brain of the mThy1-hSNCA mice (Fig. 7F). We further validated the protective effect of CS2 using immunostaining. Comparing with that in the WT littermates, the fluorescence intensity of ClpP was significantly reduced in the TH+ DA neurons in the SN of the mThy1-hSNCA mice, which was rescued upon CS2 treatment (Fig. 7G and H). Considering that neuroinflammation is a featured pathological event triggered by αSyn, we then stained IBA1 and GFAP, the markers of inflammatory-associated microglia and astrocytes, in the mid-brain. Comparing to the WT littermates, the fluorescence intensity of IBA1 and GFAP were significantly escalated in the mThy1-hSNCA mice (Fig. 7I and J), suggesting the upregulation of immune responses. While had no observed effect on the WT mice, CS2 treatment attenuated the intensity of IBA1 and GFAP signal in the mid-brain of the mThy1-hSNCA mice (Fig. 7H and I). Taken together, these findings indicate that CS2 treatment mitigated αSyn-related neuropathology and neuroinflammation *in vivo*.

**Figure 7.**
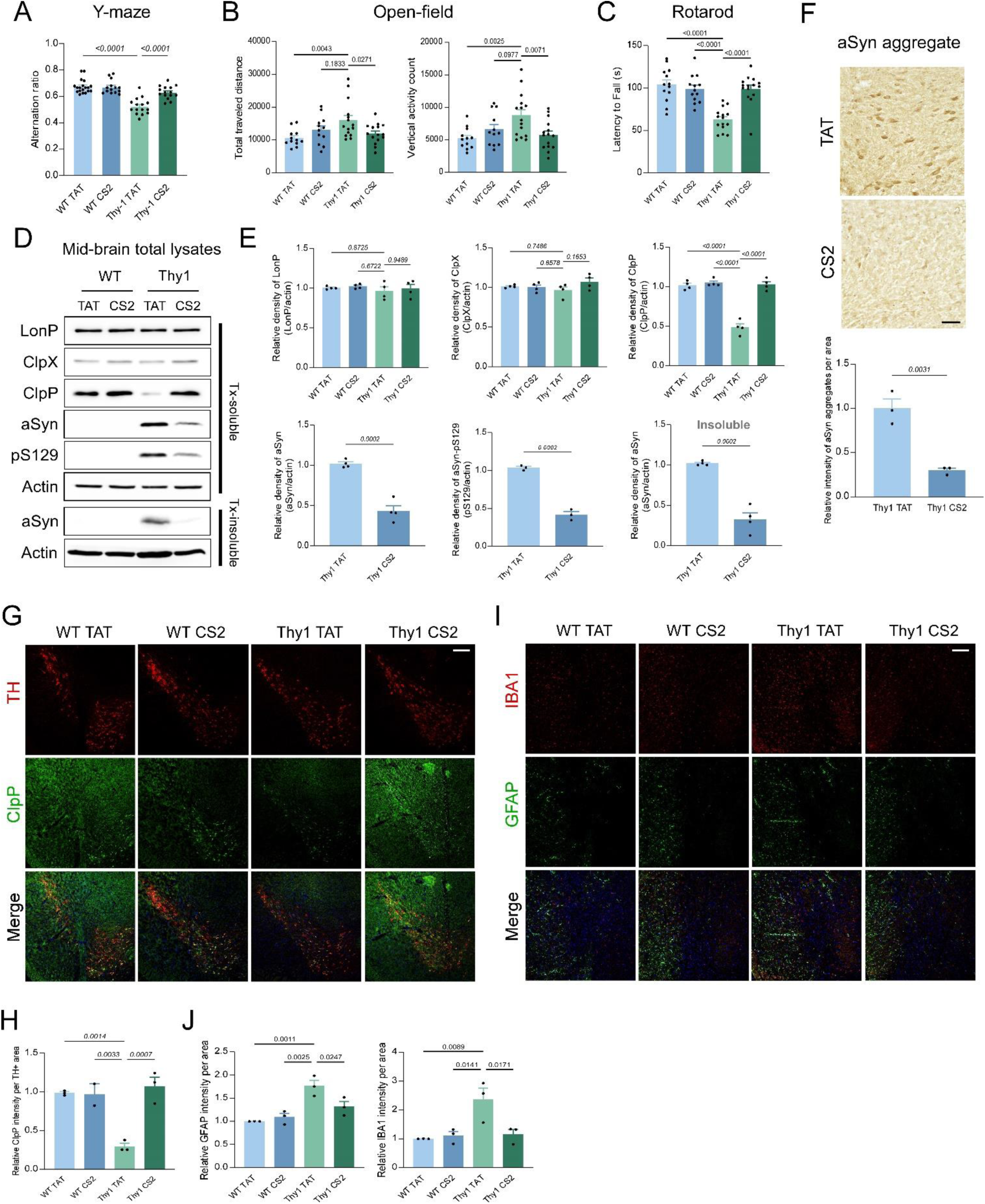
CS2 treatment is protective in mThy1-hSNCA PD mice. TAT or CS2 peptides (3mg/kg/day) were administered via osmotic pump in mThy1-hSNCA mice and their WT littermates starting from 3-month-old till 10-month-old. (A) Y-maze test was conducted at age of 6-month. Histogram: alternation ratio. N=15 for each group. (B) 24h open-field test was conducted at age of 8-month. Histogram: total travel distance and vertical activity. n=13 for WT mice, n=15 for mThy1-hSNCA mice. (C) Rotarod acceleration test was conducted at age of 10-month. Histogram: latency to fall. n=13 for WT mice, n=15 for mThy1-hSNCA mice. (D) Triton-X100 soluble and insoluble lysates were harvested from the mid-brain of 10-month WT or mThy1-hSNCA mice and subjected for western blot analyses. Shown are the representative blots, n=4 mice for each group. (E) Histograms: relative expression of the indicated protein in the soluble or insoluble fractions. (F) Immunofluorescence staining of TH (red) and ClpP (green) on the brain coronal sections from TAT or CS2 treated mThy1-hSNCA and their WT littermates. Shown are the representative images of 3 independent experiments. Scale bar=180µm. (G) Histogram: relative ClpP fluorescence intensity per TH+ area. (H) Immunofluorescence staining of Iba1 (red) and GFAP (green) in substantia nigra on the brain coronal sections from TAT or CS2 treated mThy1-hSNCA and their WT littermates. Shown are the representative images of 3 independent experiments. Scale bar=180µm. (I) Histogram: relative Iba or GFAP fluorescence intensity per 4.2 mm^2^. (J) Immunohistochemistry staining of α-Syn aggregates in substantia nigra by using specific anti-α-Syn aggregates antibody (5G4) on the coronal sections from TAT or CS2 treated mThy1-hSNCA and their WT littermates. Shown are the representative images of 3 independent experiments. Scale bar=40µm. Histogram: relative intensity of α-Syn aggregates per 1 mm^2^. All data are expressed as the mean ± SEM. Data are compared with one-way ANOVA with Tukey’s *post-hoc* test or unpaired student *t*-test.

## DISCUSSION

In this study, by focusing on the pathological interaction between α-Syn and ClpP, we identified several key findings. First, we discovered that ClpP acts as a negative regulator of α-Syn aggregation and propagation, likely by stabilizing α-Syn tetramers. We also demonstrated that the NAC domain of α-Syn is required for its interaction with ClpP, which is associated with the downregulation of ClpP expression and its enzymatic activity. Furthermore, our rationally designed peptide, CS2, efficiently blocked the α-Syn/ClpP interaction and inhibited α-Syn aggregation in vitro. CS2 treatment substantially restored ClpP expression and ameliorated α-Syn-induced neurotoxicity *ex vivo*. Moreover, sustained administration of CS2 significantly mitigated α-Syn-related neuropathology and improved cognitive and motor deficits in mThy1-hSNCA mice. Overall, these findings indicate that loss of ClpP exacerbates α-Syn aggregation and propagation—critical components of PD pathophysiology—and that disrupting the α-Syn/ClpP interaction may represent a promising therapeutic strategy for neurodegenerative disorders characterized by α-Syn aggregation.

Emerging evidence suggests that α-Syn naturally exists as a helically folded tetramer, which is resistant to aggregation^14^. Disrupting α-Syn tetramerization leads to the accumulation of unfolded monomers, facilitating and accelerating α-Syn aggregation and propagation in PD pathogenesis^14^. Thus, α-Syn tetramerization plays a critical role in controlling the formation of toxic α-Syn species, and understanding its regulation could provide key insights into PD etiology. Here, we found that ClpP deficiency is sufficient to shift α-Syn from its tetrameric to monomeric form. Conversely, ClpP overexpression significantly increased the α-Syn tetramer-to-monomer ratio, suggesting that ClpP may positively regulate native α-Syn tetramers. However, whether ClpP modulates α-Syn through direct interaction or indirectly via mitochondrial pathways remains to be determined. Notably, overexpression of the proteolytically inactive ClpP-S153A mutant significantly reduced α-Syn tetramer levels. Given that ClpP-S153A upregulation leads to the accumulation of mitochondrial proteomes, we postulate that ClpP-mediated mitochondrial proteostasis may be crucial for maintaining the α-Syn tetramer pool. Further research is needed to elucidate the precise mechanisms by which ClpP regulates α-Syn tetramerization. It would also be interesting to investigate whether ClpP influences α-Syn aggregation and propagation primarily by stabilizing its tetrameric form or through alternative mechanisms. Such insights could significantly enhance our understanding of α-Syn regulation in PD pathogenesis.

Current treatments for PD, including levodopa-like agents, dopamine agonists, and surgical interventions such as deep brain stimulation, primarily alleviate motor symptoms without altering disease progression^30, 31^. Over time, these therapies become less effective and often lead to significant side effects, including dyskinesia. Moreover, effective treatments for the debilitating non-motor symptoms of PD are lacking, and no existing therapy can halt or slow the neurodegenerative process underlying the disease. Given that αSyn aggregation and mitochondrial damage are pivotal pathological features of PD, therapeutics that target the vicious cycle between mitochondrial dysfunction and αSyn toxicity may offer promising treatment strategies. Our findings demonstrate that CS2 treatment is protective in models of αSyn-induced neurotoxicity, providing proof of concept that peptide inhibitors disrupting the mitochondria–αSyn interaction may serve as a potential therapeutic strategy for PD. Notably, beyond blocking the interaction between ClpP and αSyn, CS2 directly binds to αSyn and modulates its aggregation. The CS2 peptide sequence is derived from a region of ClpP that aligns with the NAC domain of αSyn. Intriguingly, the proposed CS2 binding pocket contains several critical residues that are essential for the formation of the unfolded core of αSyn fibrils, which may explain why CS2 can directly suppress αSyn aggregation in our models. In the near future, we plan to investigate the “one stone, two birds” effect of CS2 in models of αSyn toxicity—a dual-target approach that may prove more advantageous than conventional therapies targeting a single mechanism. To fully evaluate the therapeutic potential of CS2, we will conduct *in vivo* pharmacokinetic and pharmacodynamic analyses.

## METHODS

### Antibodies and reagents

Protein phosphatase inhibitor and protease inhibitor cocktails were purchased from MilliporeSigma. For western blot, antibodies against ClpX (ab168338, 1:2000), ClpP (ab124822, 1:1000), α-Synuclein (ab138501, 1:3000), and pS129-α-synuclein (ab168381, 1:1000) were from Abcam. Antibodies against GFP (sc-9996, 1:1000), c-Myc (sc-40, 1:1000), α-Synuclein (sc-12767, 1:1000), were from Santa Cruz Biotechnology. Antibodies against Flag (F1804. 1:5000), β-Actin (A1978, 1:10000) were from MilliporeSigma. Antibodies against LONP1 (15440-1-AP, 1:3000) was from Proteintech.

For immunostaining, antibodies against TH (MAB318, 1:1000), NeuN (ABN90, 1:500), and α-Synuclein aggregates (clone 5G4) (MABN389, 1:500) were from MilliporeSigma. Antibody against TOM20 (11802-1-AP, 1:2000) was from Proteintech. Antibodies against Calnexin (ab22595), and pS129-α-synuclein (EP1536Y, 1:1000) were from Abcam. Antibodies against α-synuclein (Syn204) (2647, 1:500), and MAP2 (4542, 1:1000) were from Cell Signaling Technology. Antibody against Tuj1 (801201, 1:1000) was from Biologend. Antibody against Synapsin1 (106104, 1:500) was from SYSY Synaptic System. Antibody against PSD95 (MA1-045, 1:500) was from ThermoFisher Scientific. Antibody against ClpP (NBP1-89557, 1:200) was from Novus Biologicals. Antibody against TH (TYH0020, 1:1000) was from avesLab.

### Cell culture

SH-SY5Y cells were cultured in Dulbecco’s Modified Eagle Medium (DMEM)/F12 (1:1) supplemented with 10% (v/v) fetal bovine serum (FBS) and 1% (v/v) antibiotics (100 µg/ml penicillin and 100 µg/ml streptomycin) at 37℃ in a 5% CO_2_ incubator. HEK293T cells were cultured in DMEM supplemented with 10% (v/v) FBS and 1% (v/v) antibiotics.

### Preparation of total cell lysates

Cells were washed with 1X phosphate-buffered saline (PBS) and then incubated in total lysis buffer (10 mM HEPE-NaOH [pH 7.8], 150 mM NaCl, 1 mM ethylene glycol-bis [β-aminoethyl ether]-*N*,*N*,*N’*,*N’*-tetraacetic acid [EGTA], 1% Triton X-100, protease inhibitors, and phosphatase inhibitors) for 30 min on ice. Samples were collected and centrifuged at 12,000 rotations per minute for 10 min at 4°C. The supernatants were saved as total lysates.

### Western blot

Protein concentrations were measured using the Bradford assay. Proteins (30 μg) were then resuspended in Laemmni buffer, loaded on sodium dodecyl sulfate-polyacrylamide gel electrophoresis gels, and transferred to nitrocellulose membranes. The membranes were probed with the indicated antibodies and visualized by electrochemiluminescence.

### Constructs and transfection

EGFP-tagged WT αSyn (40822), Venus-N-αSyn (89470), and αSyn-C-Venus (89471) were purchased from Addgene. Myc-tagged αSyn-WT and –A53T, Myc-tagged ClpP, and Flag-tagged WT and S153A ClpP were constructed as previously described^12^. Cells were transfected with *Trans*IT-2020 (Mirus Bio, LLC) following the manufacturer’s protocol.

### Peptides treatment in primary neuronal model of alpha-synuclein PFF-induced toxicity

Primary cortical neurons were isolated from E18 mouse cortex and cultured using complete culturing kit (KTC57ECX, TransnetYX Tissue). Briefly, 100,000 cells were seeded on poly-D-lysine coated glass coverslips (PDL15mm, TransnetYX Tissue) and cultured in NbActiv1 medium (NB1, TransnetYX Tissue). Neurons were treated with sonicated human α-Syn-PFF (SPR-322, StressMarq Biosciences) starting on post-seeding Day 7 till Day 14. Starting from post-seeding Day 8, peptide TAT or CS2 was added daily at 1μM. Neurons were fixed on Day 14 and subjected for immunostaining.

### Peptides administration in mThy1-hSNCA mice

All animal experiments were conducted in accordance with protocols approved by the Institutional Animal Care and Use Committee of Case Western Reserve University and were performed based on the National Institutes of Health Guide for the Care and Use of Laboratory Animals. Sufficient procedures were employed for reduction of pain or discomfort of subjects during the experiments.

C57BL/6N-Tg(Thy1-SNCA)15Mjff/J (JAX: 017682) breeders (C57Bl/6NJ genetic background) were purchased from Jackson Laboratories. The mice were mated, bred, and genotyped in the animal facility of Case Western Reserve University. All mice were maintained at a 12-hour light/dark cycle (on 6 am, off 6 pm). Control peptide TAT and CS2 peptide (Cat#: P2 YA-18, Lot#: P190717-MJ738691) were synthesized at Ontores (Hangzhou, China). Their purities were assessed as >90% by mass spectrometry. Lyophilized peptides were dissolved in sterile water and stored at -80°C until use. All randomization and peptide treatments in mThy1-hSNCA mice were prepared by an experimenter not associated with the behavioral and neuropathology analysis. mThy1-hSNCA transgenic mice and their age-matched and sex-balanced WT littermates were implanted with a 28-day osmotic pump (Alzet, Cupertino CA, Model 2004) containing either TAT control peptide or DA1 peptide, which delivered the peptides at a rate of 1mg/kg/day. The pump was replaced once every four weeks starting from age of 16-week. All mice were sacrificed at age of 40-week for neuropathology assessment.

### Behavioral analysis

All behavioral analyses were conducted by an experimenter who was blinded to the genotypes and treatment groups. All mice were subjected to a series of behavioral measurements to monitor locomotor activity (open field test), spontaneous spatial working memory (Y-maze test), and motor coordination (Rotorod test).

### Y-maze test

On the test day, mice at 6-month-old were brought to the testing room one hour before performing the Y-maze test to allow habituation. The mice were placed in the middle of the Y-maze and allowed to explore the three arms for 6 min. During exploration, the arm entries were recorded. The equipment was cleaned after every test to avoid odor disturbance. Spontaneous alternation was defined as a successive entry into three different arms on overlapping triplet sets.

### Open field test

The locomotor activity of all experimental mice was assessed in an open field at 8-month-old. Briefly, the mice were placed in the center of an activity chamber (Omnitech Electronics) and allowed to explore the chamber while being tracked using an automated infrared tracking system (Vertax, Omnitech Electronics). A 24-h locomotor activity analysis was performed. Both horizontal and vertical activity/movements were recorded.

### Rotarod test

Motor coordination was assessed in TAT or CS2 treated mThy1-hSNCA transgenic mice and their age-matched littermates at age of 8-month, by measuring latency on a rotarod in an accelerated program for 300 s at 10-month-old. Body weights of mThy1-hSNCA mice and WT littermates were recorded throughout the study period.

### Peptides treatment in human iPSC-derived neurons

PD iPSC lines (αSyn A53T, NN0004337) and its isogenic control line (NN0004344) were obtained from RUCDR Infinite Biologics. The iPS cells were differentiated into DA neuron-enriched neuronal culture with the protocol described previously ^12, 32, 33^. Briefly, iPS cell colonies were disassociated with accutase (Invitrogen), plated onto 6-well plates pre-coated with 2.5% Matrigel (BD Biosciences) and allowed to reach 80% confluence in mTeSR medium (Stem Cell Technology). For the first 10 days, cells were treated with SB431542 (10 uM; Tocris Bioscience) and Noggin (100 ng/ml) in Neural Media (NM) with FGF2 (20 ng/μl) and EGF (20 ng/μl). NM media contained: Neurobasal and DMEM/F12 (1:1), B-27 Supplement Minus Vitamin A (50×, Invitrogen), N2 Supplement (100X, Invitrogen), GlutaMAX (Invitrogen, 100×), 100 units/ml penicillin and 100 μg/ml streptomycin (Fisher); for the next 4 days, cells were treated with human recombinant Sonic hedgehog (SHH, 200 ng/ml) in neuronal differentiation medium. Neuronal differentiation medium contained Neurobasal and DMEM/F12 (1:3), B27, N2, GlutaMax and PS. In the following 3 days, cells were switched to BDNF (20 ng/ml), ascorbic acid (200 uM, Sigma-Alderich), SHH (200 ng/ml), and FGF8b (100 ng/ml) in neuronal differentiation medium. Thereafter, cells were treated with BDNF, ascorbic acid, GDNF (10 ng/ml), TGF-b (1 ng/ml), and cAMP (500 uM, Sigma-Aldrich). All growth factors were purchased from Pepro Tech (Rocky Hill, NJ, USA). After 20 days of induction, neurons were treated daily with TAT or CS2 peptides at 1µM and were fixed for immunostaining at 25 days of induction. The imaging was observed by confocal microscope (Fluoview FV3000, Olympus).

### Intact cell crosslinking to detect alpha-synuclein tetramer

*After wash with PBS (pH 8.0), cell pellets were incubated in 1mM DSG solved in PBS for 40 min at* 37 °C with 650 rpm shaking on Eppendorf ThermoMixer C. The reaction was then quenched with 50mM Tris pH 7.4 for 15 min at room temperature. The cells were then lysed by ultrasonication for 15s at 20% amplitude (Branson Digital Sonifier). Cells lysates were then centrifuged at 20,000g at 4°C for 40 min. The supernatant was then collected for western blot analysis. Alpha-synuclein monomer and oligomers can be detected using anti-α-Synuclein (sc-12767, 1:1000) antibody from Santa cruz Biotechnology.

### Alpha-synuclein ThT assays

*Alpha-synuclein monomer (5µM) and PFF (1µM) were incubated with recombinant ClpP (5/10µM) and Thioflavin T (ThT) (1µM) in 100µl PBS (pH 7.4) at 37*°C in a shaking incubator (500 rp,m)*. ThT fluorescence was measured (excitation 450 nm, emission 485 nm) every 15 min up to 48h*.

### Immunofluorescence

Cells were grown on coverslips, fixed with 4% paraformaldehyde for 20 min at room temperature, permeabilized with 0.1% Triton X-100 in PBS, and blocked with 2% normal goat serum. The cells were incubated with the indicated primary antibodies overnight at 4℃. After washing with PBS, the cells were incubated with Alexa Fluor 488/568 or 405/568 secondary antibody (1:500; Thermo Fisher Scientific) for 2 h at room temperature. The nuclei were counterstained with DAPI (1:10000; Sigma-Aldrich). Images of the staining were acquired using a Fluoview FV3000 confocal microscope (Olympus).

For immunofluorescence staining of mouse brain sections, mice were deeply anesthetized and transcardially perfused with 4% paraformaldehyde in PBS. Brain sections were permeabilized with 0.2% Triton X-100 in TBS-T buffer, followed by blocking with 5% normal goat serum. The brain sections were incubated with the indicated primary antibodies overnight at 4°C and then stained with secondary antibodies. Images of the staining were acquired using a Fluoview FV3000 confocal microscope (Olympus).

All quantification of immunostaining was performed using ImageJ software. The same image exposure times and threshold settings were used for all sections from all the experimental groups. Quantitation was performed blinded to the experimental groups.

### Immunohistochemistry

Frozen brain sections (14 μm, coronal) were staining for alpha-synuclein aggregates (MABN389, clone 5G4, Millipore) using the IHC Select HRP/DAB kit (Millipore). Quantification of the alpha-synuclein immunostaining was conducted using NIH Fiji ImageJ software. The same image exposure times and threshold settings were used for all sections from all treatment groups. Quantitation was performed blinded to the experimental groups.

### ClpP peptidase activity in vitro

ClpP peptidase activity in vitro was measured as previously described. Briefly, human recombinant ClpP (10 µM, obtained from Dr. Aaron Schimmer, Princes Margartet Cancer Centre, Canada) were incubated in the reaction buffer (50mM Tris PH 8.0, 200mM KCl, 1mM DTT, 2mM ATP) under 37°C for 10 min. For co-incubation with α-synuclein, recombinant ClpP and WT or C-terminal truncated or NAC-domain truncated recombinant αSyn (S-1001-1, S-1012-1, S-1015-1, rPeptide) were pre-incubated for 30 min under room temperature. Fluorescent substrate of ClpP, ac-WLA-AMC (50 µM), were then added in the reaction buffer. The fluorescence signal was read using TECAN infinite M1000 up for 30 min at excitation/emission wavelength 345/445. ClpP peptidase activity were determined as the slope of the regression line.

### Co-immunoprecipitation

Cells were lysed in a total cell lysate buffer (50mM Tris-HCl, pH 7.5, 150mM NaCl, 1% Triton X-100, and protease inhibitor). Total lysates were incubated with the indicated antibodies overnight at 4°C followed by the addition of protein A/G beads for 2 h at 4°C.

Various recombinant α-Synuclein and Clpp (500 ng) were incubated in *in vitro* interaction buffer (20mM Tris-HCl pH 7.5, 100mM KCl, 2mM MgCl_2_ and 0.1% Triton-X100) for 30 minutes at room temperature, and then incubated with indicated antibodies overnight at 4°C followed by the addition of protein A/G beads for 2h. Immunoprecipitates were washed four times with cell lysate buffer/in vitro interaction buffer in the presence of 0.1% Triton X-100 and were analyzed by western blot analysis.

### Mitochondrial ROS measurement

Cells cultured on coverslips or 24-well plates were washed with DPBS and then incubated with 5 µM MitoSOX^TM^ Red (Invitrogen, M36008), a mitochondrial superoxide indicator, for 10 min at 37 °C. For cells cultured on coverslips, the images were visualized by microscope, and quantification of images was then carried out using NIH ImageJ software. At least 100 cells per group were counted in the analysis

### Cell death measurement by LDH

SHSY5Y cells were treated with different doses of TAT/CS2 peptides for 3 days, and cell death was determined using the Cytotoxicity Detection Kit (LDH), according to the manufacturer’s protocol (Roche, REF 11 644 793 001).

### Alpha-synuclein ELISA

After one day incubation with doxycycline (1µM), alpha-synuclein tet-on inducible expression SHSY5Y cells were treated with TAT or CS2 peptides at 10 µM for another two days. The level of alpha-synuclein monomer released into culture medium was measured using LEGEND MAX human a-Syn ELISA kit (448607, Biolegend). The level of extracellular and intracellular alpha-synuclein oligomers was measured using human alpha synuclein oligomer ELISA kit (MBS730762, Mybioresource).

## ACKNOWLEDGMENTS

This study was supported by grants from the US National Institutes of Health (R01AG065240, R01NS115903, R01AG076051 and RF1AG074346 to X.Q.); Vinney Scholar Award for Alzheimer’s disease to X.Q.

## AUTHOR CONTRIBUTIONS

D.H. designed experiments, performed experiments and analyses, and drafted the manuscript. X.S. maintained the mouse line. X.Q. conceived, designed, and supervised all the studies and edited the manuscript.

---

**Figure S1.**
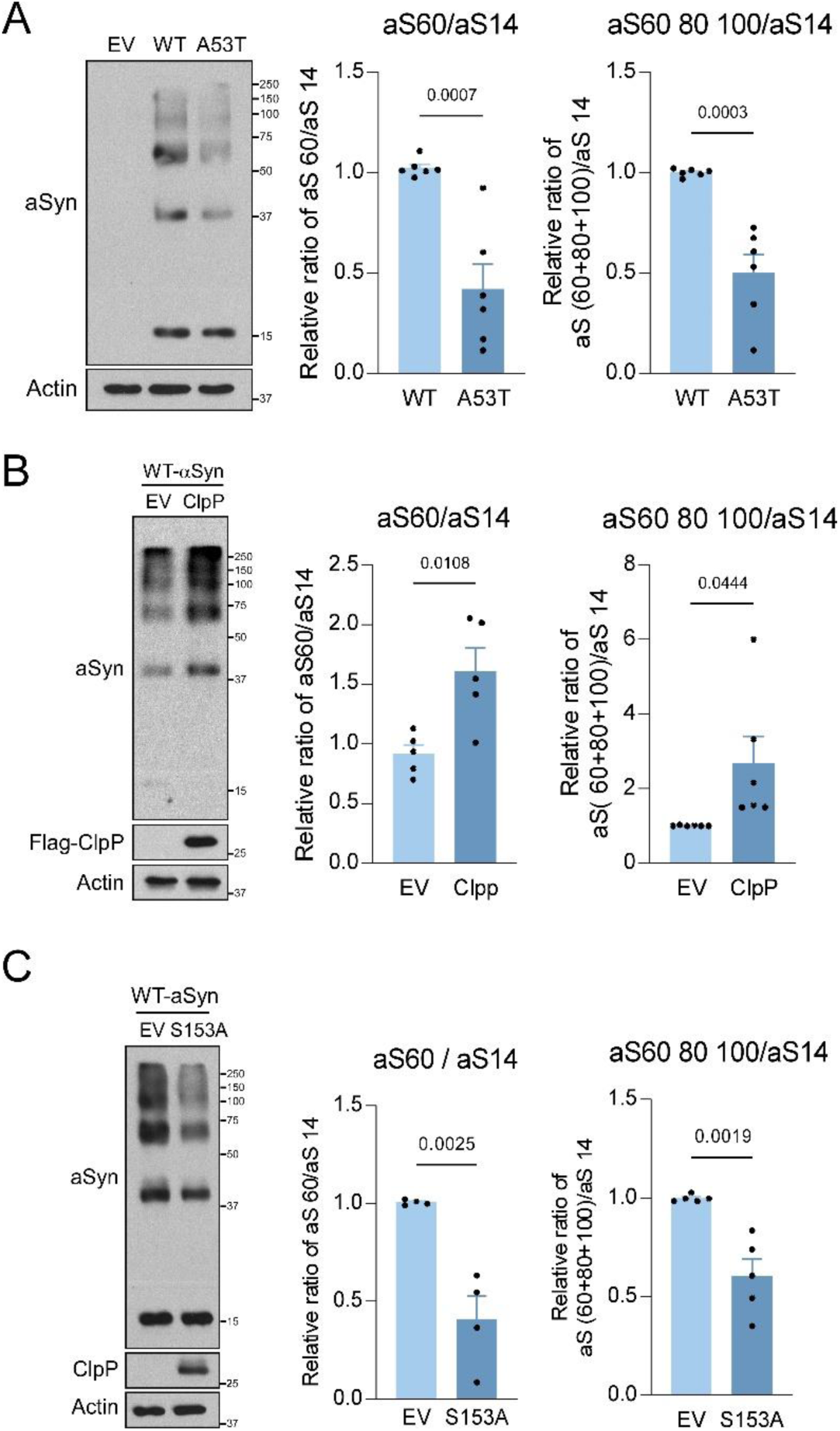
ClpP modulates α-Syn tetramers. (A) HEK293T cells overexpressing control vector (EV), WT-α-Syn, or A53T-α-Syn were subjected for intact cell crosslinking by DSG (see methods), followed by western blot analyses. Shown are the representative blots of 6 independent experiments. Histogram: relative ratio of α-Syn tetramer to monomer, and α-Syn oligomers to monomer. HEK293T cells overexpressing (B) Myc-tagged WT-α-Syn or (C) Myc-tagged A53T-α-Syn, and control vector (EV) or Flag-tagged ClpP, were subjected for intact cell crosslinking by DSG, and followed by western blot analyses. Shown are the representative blots of at least 4 independent experiments. Histogram: relative ratio of α-Syn tetramer to monomer, and α-Syn oligomers to monomer. (D) HEK293T cells overexpressing WT-α-Syn and EV or S153A mutant ClpP were subjected for intact cell crosslinking by DSG and followed by western blot analyses. Shown are the representative blots of 4 independent experiments. Histogram: relative ratio of α-Syn tetramer to monomer, and α-Syn oligomers to monomer. All data are expressed as the mean ± SEM. Data are compared with unpaired student *t*-test.

**Figure S2.**
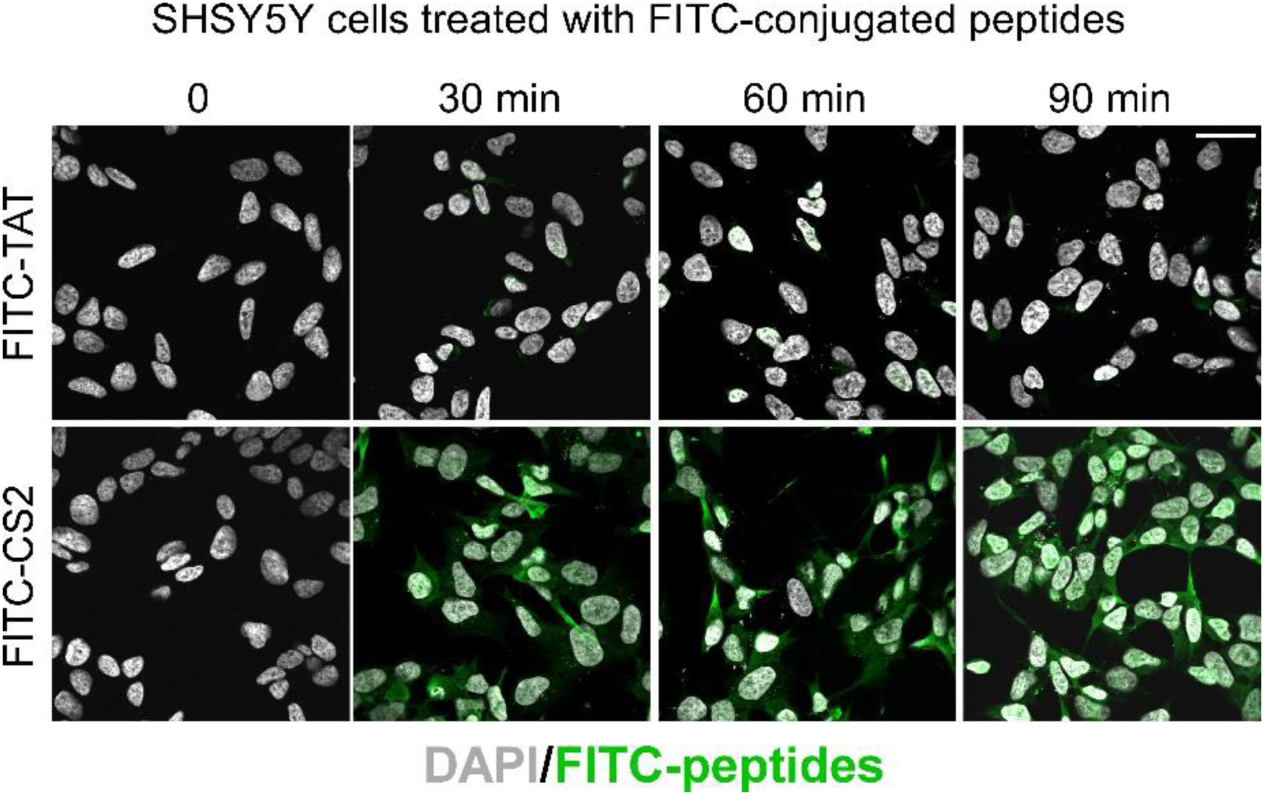
CS2 peptide enters cell in 30 min. SH-SY5Y cells were treated with FITC-conjugated TAT or CS2 peptides (10 μM). Cells were fixed at different time points as indicated and subjected for confocal imaging. Shown are the representative images. Scale bar=30µm.

**Figure S3.**
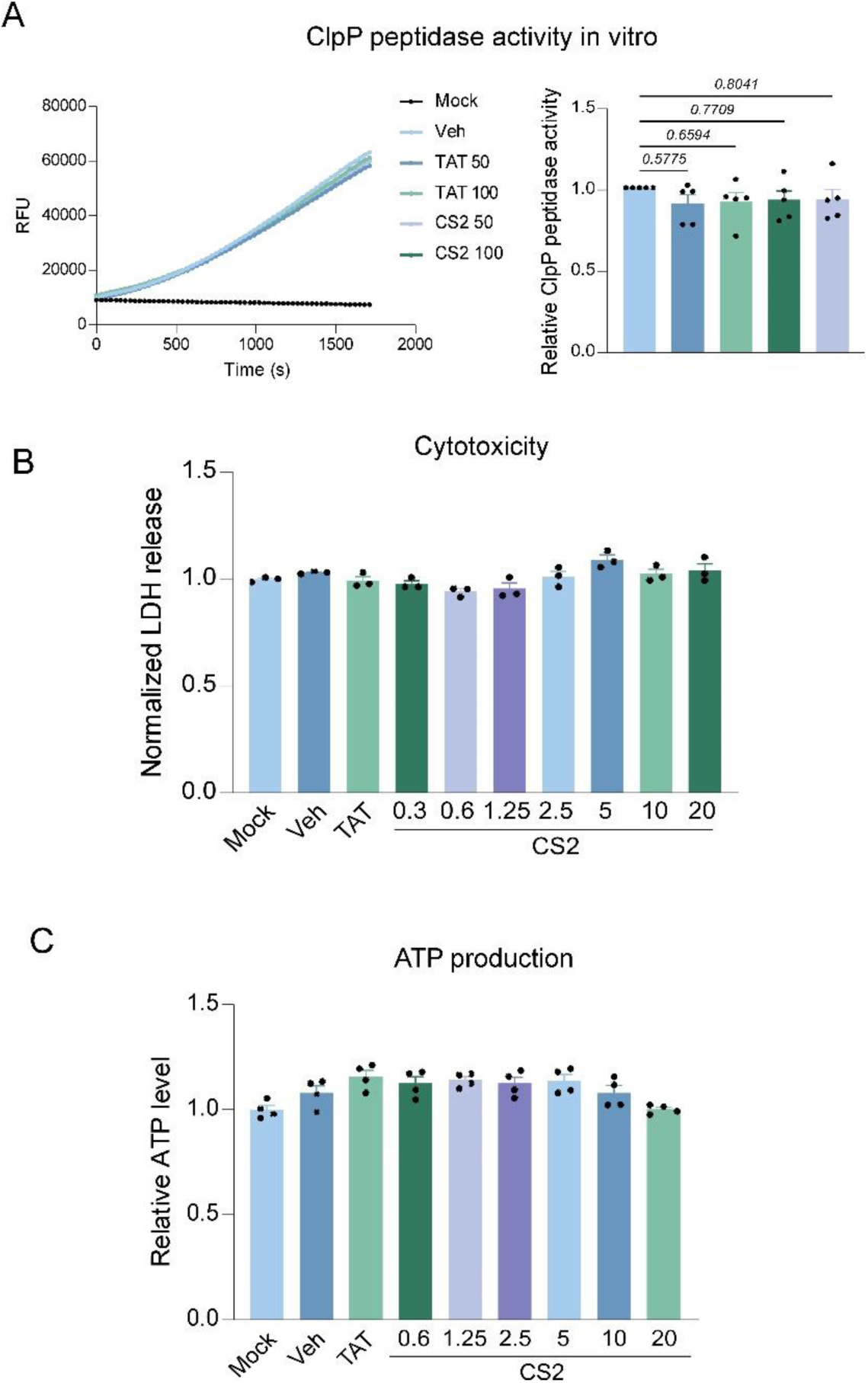
CS2 has minimal toxicity in cells. (A) In vitro ClpP peptidase activity was measured in presence of TAT or CS2 peptides (50/100 µM). The fluorescence intensity of ac-WLA-AMC (50 µM), a fluorogenic substrate of ClpP, was measured up to 30 min immediately after the addition. Shown are the representative RFU measurement of 5 independent experiments. Histogram: relative ClpP peptidase activity. SHSY5Y cells were treated with TAT or CS2 peptides with the indicated doses for 48h. (B) Cell death was measured by LDH assay. N=3 independent experiments. (C) Cell ATP level was measured and quantified as shown in histogram. N=4 independent experiments. All data are expressed as the mean ± SEM. Data are compared with one-way ANOVA with Tukey’s *post-hoc* test.

